# Large language models and network medicine to identify drug combinations for estrogen receptor-positive breast cancer: a computational study

**DOI:** 10.64898/2026.09.14.728753

**Authors:** Ahmed Abdeen Hamed, Tamer E. Fandy, Luis M. Rocha

**Affiliations:** Department of Biochemistry, University of Nebraska–Lincoln, Lincoln, NE 68588, USA; School of Systems Science and Industrial Engineering, Binghamton University, Binghamton, NY 13902, USA; Paul L. Foster School of Medicine, Texas Tech University Health Sciences Center El Paso, El Paso, TX 79905, USA; Universidade Catolica Portuguesa, Catolica Biomedical Research Centre, Lisbon, Portugal

## Abstract

**Background:** Repurposed drug combinations could offer new treatments for estrogen receptor-positive (ER^+^) breast cancer, but the evidence to choose them is scattered across millions of publications and trial records. We assessed whether large language models (LLMs) can assemble this evidence reliably at scale.

**Methods:** Six open-weight LLMs extracted drug–target relationships from 587,256 PubMed abstracts; relationships reported by at least two models were used to build a consensus network. A seventh model extracted drug combinations from 2,496 ClinicalTrials.gov descriptions by majority vote across three prompts (self-consistency). Drug and target names were mapped to ChEBI and HGNC; mapping accuracy was measured on random samples by majority vote of three additional LLMs. Complementary drug pairs on the KEGG estrogen signaling pathway (each drug reaching branches the other does not; synergy was not assessed) were identified. Literature evidence was cross-checked by retrieval-augmented extraction; pairs were ranked by robust rank aggregation and evaluated against trial precedent.

**Findings:** The final consensus network contained 760,743 drug–target relationships, and 2,187 drugs reached 35 of 63 estrogen signaling pathway genes. Accuracy based on a fresh validation sample was 91.0% (95% CI 86.2–94.2) for drugs and 94.0% (89.8–96.5) for targets. Retrieval-augmented and bulk extraction agreed on pathway-level pairs (odds ratio 10.9, *p <* 0.0001). Of 81,207 complementary pairs, 94 had trial precedent. Among the top 100 ranked pairs, 10.6% had trial precedent, compared with 0.12% of all pairs, but only through trial-seeded retrieval evidence; pathway coverage and model consensus alone recovered none.

**Interpretation:** Consensus ensembles of LLMs can extract mechanistic evidence for combinatorial repurposing at a scale beyond manual curation, and mapping to curated references with measured accuracy shows that the evidence is usable. It can be used to prioritize combinations for expert review, but it does not by itself predict clinical success.

**Funding:** SUNY AI Platform, European Commission, Fundação para a Ciência e a Tecnologia.

**Research in context**

Evidence before this study
We searched PubMed on Nov 12, 2026, for studies published from Jan 1, 2018, to Nov 12, 2026, with no language restrictions, using terms for large language models (LLMs) and drug repurposing in titles and abstracts: (“large language model*” OR “LLM” OR “LLMs” OR “GPT” OR “ChatGPT” OR “generative AI”) AND (“drug repurposing” OR “drug repositioning”). We found 90 articles and reviewed the 25 most relevant. These studies mainly used LLMs to suggest individual drugs for diseases, find links between drugs and genes or diseases in published research, or support AI tools. None looked at using LLMs to find combinations of drugs for breast cancer. Research on using networks of biological information to find drugs that could be repurposed for breast cancer is more established (21 articles in the same period), but it has generally relied on information already collected in databases rather than evidence gathered at scale from published research. An earlier AI-assisted search of Scopus (LeapSpace, May 25, 2026; query not retained) found 31 articles on computer-based approaches to finding new uses for drugs, mostly for breast cancer. One proposed drug combinations for luminal A disease. We did not formally assess the risk of bias in these studies; most described methods rather than evaluating clinical outcomes. We found no study that used LLMs to find and check evidence about how drug combinations might work in ER^+^ breast cancer.

Added value of this study
Six open-weight LLMs read 587,256 PubMed abstracts, and another model read 2,496 clinical trial descriptions. We used agreement between models and across prompts to reduce reliance on any single model’s answers. Drug and target names were matched to reference databases (ChEBI and HGNC) and checked by a separate group of models. The checks found that 91.0% of drug mappings and 94.0% of target mappings were correct. Two approaches to reviewing the literature—reading abstracts in bulk and searching for evidence related to specific drugs—found overlapping links between drugs and targets in the estrogen signaling pathway (odds ratio 10.9). However, pathway coverage and model agreement alone did not rank combinations that had been tested in clinical trials near the top of the list.

Implications of all the available evidence
LLMs can help gather evidence about how drugs might work together on a scale that would be difficult to manage by hand. This evidence can help organize drug combinations for expert review, but it cannot show whether a combination will help patients. That will require evidence independent of existing trial records and testing in experiments. The approach uses open-weight models and openly available code, and could be adapted to other diseases and to full-text articles.

## Introduction

Drug repurposing, the use of approved or clinically tested drugs for new indications, can shorten the path to new treatments because each drug’s pharmacology and safety are already partly known ^1^. Combinations of repurposed drugs extend this idea: two drugs acting on different parts of a disease’s signaling network may achieve what neither achieves alone ^2^. ER^+^ breast cancer, the most common subtype, is a natural setting for this approach, because its growth depends on estrogen receptor signaling and on the growth-factor pathways that interact with it ^3–5^. The difficulty is not a lack of candidate drugs but a lack of a systematic way to assemble the evidence on which combinations are worth testing.

Network medicine offers the scaffold for that assembly. Representing genes, drugs, and diseases as nodes in interaction networks reveals relationships that no single target shows ^6,7^, and the proximity of a drug’s targets to a disease module has been used to propose new indications ^8,9^. The way two drugs’ targets sit relative to a disease module has been used to predict effective combinations ^10^. Network-based studies have identified repurposable drugs for breast cancer subtypes from transcriptomic, interactomic, and pathway data ^5,11,12^, and one proposed drug combinations for luminal A breast cancer from expression data ^13^. These approaches, however, depend largely on curated databases and experimental profiles, which lag behind the literature. Our earlier work showed that networks built directly from the biomedical literature and from clinical trial records can surface repurposing candidates ^14,15^, but extracting relationships from millions of documents by hand or with rule-based tools does not scale.

Large language models (LLMs) can read at that scale, and they can be instructed in natural language to extract drug–target relationships from text. Their outputs, however, vary across models and tasks and can include relationships that no document states ^16–18^. Grounding LLM outputs in structured knowledge, such as knowledge graphs or retrieved evidence, is an emerging strategy for making them reliable ^19–21^. We previously combined network medicine with generative AI in a breast cancer case study ^22^; what remains open is whether LLM extraction can be made trustworthy at the scale of the whole literature, and what such evidence can and cannot support.

In this study, six open-weight LLMs read 587,256 PubMed abstracts and extracted drug–target relationships, keeping only those reported by at least two models; a further model extracted drug combinations from 2,496 clinical trial descriptions by self-consistency. We mapped drug and target names to curated references (ChEBI and HGNC), measured the accuracy of that mapping with an independent panel of models, and grounded the evidence in the KEGG estrogen signaling pathway. We then enumerated drug pairs that act on complementary branches of the pathway, compared them with clinical trial records, which served as the benchmark for the ranking, checked the literature evidence against a second, retrieval-based extraction, and ranked the pairs for expert review. Our aim is to show how LLMs can accelerate the assembly of mechanistic evidence for combinatorial repurposing, and to report plainly what that evidence supports.

## Methods

We designed the study in five steps (Figure 1). Six open-weight LLMs extracted drug–target relationships from PubMed abstracts, and a seventh extracted drug combinations from clinical trial records. Drug and target names were mapped to curated references, and the mapping was validated. The evidence was overlaid on the Kyoto Encyclopedia of Genes and Genomes (KEGG) estrogen signaling pathway, complementary drug pairs were enumerated, and pairs were ranked by robust rank aggregation and evaluated against trial precedent, which was not a ranking input, although two inputs derive from searches seeded with trial drugs. LLMs read the abstracts and trial records and checked a sample of name mappings; all other steps, including name mapping, pathway overlay, pair enumeration, and ranking, were rule-based. Only public data were used (PubMed, ClinicalTrials.gov, KEGG, ChEBI, and HGNC), and ethics approval was not required. Further details are in the appendix (supplementary methods).

**Figure 1:**
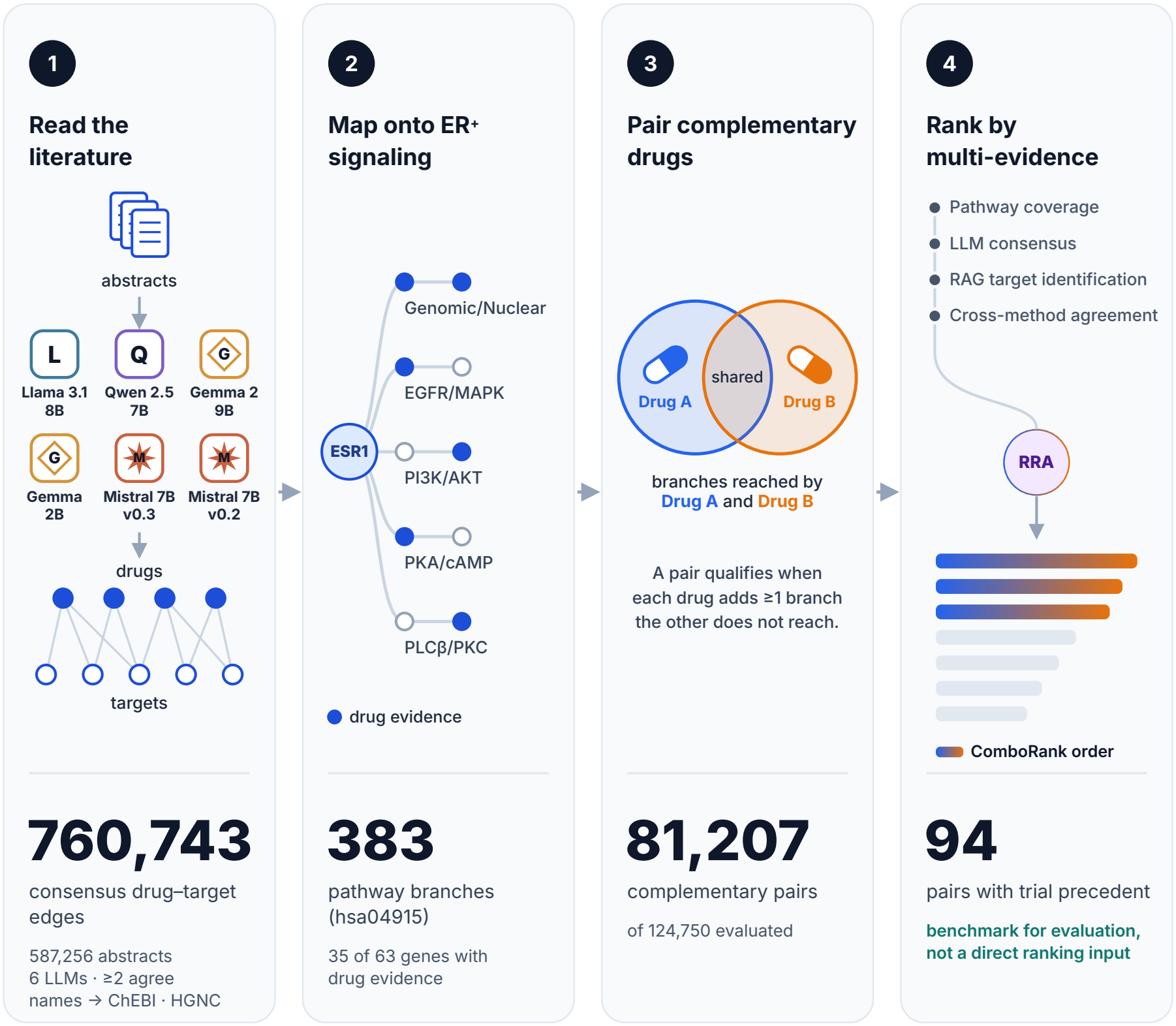
Study workflow. (1) LLM-driven evidence extraction: six LLMs read 587,256 PubMed abstracts and extract drug–target pairs; edges on which at least two LLMs agree form a consensus network of 760,743 drug–target edges, and drug and target names are mapped to ChEBI and HGNC. (2) Estrogen signaling pathway overlay: the consensus network is mapped onto the KEGG estrogen signaling pathway (hsa04915), which has 383 signaling branches; 35 of 63 pathway genes have drug evidence. (3) Drug combination discovery: a pair qualifies when each drug reaches at least one branch the other does not; 81,207 of 124,750 evaluated pairs are complementary. (4) Multi-evidence ranking: ComboRank orders pairs by Robust Rank Aggregation (RRA) of pathway coverage, LLM consensus, RAG support, and cross-method agreement; trial precedent is not a direct ranking input, although RAG support and cross-method agreement derive from searches seeded with trial drugs; the 94 pairs with trial precedent are the benchmark for evaluating it. Drawn with the assistance of Claude (Anthropic).

### LLM-driven evidence extraction

#### Literature corpus

We queried PubMed with (inhibitor*[Title/Abstract]) AND (receptor[Title/Abstract] OR kinase[Title/Abstract] OR enzyme[Title/Abstract]) and downloaded 595,122 MEDLINE records on Dec 17, 2025. Only records with an abstract were submitted for extraction, and prompts longer than 8,000 characters were skipped.

#### Drug–target extraction

Six open-weight LLMs (Llama-3.1-8B-Instruct, Qwen-2.5-7B-Instruct, Gemma-2-9B-it, Gemma-2B-it, Mistral-7B-Instruct-v0.2, and Mistral-7B-Instruct-v0.3) each read every abstract independently. A structured prompt (appendix, prompt P4) asked for each drug mentioned, its protein or molecular target, and the interaction type, returned as strict JSON. No retrieval step was used.

#### Clinical trial corpus

We retrieved 2,496 trial descriptions from ClinicalTrials.gov. Combinations were extracted with Phi-3.5-mini-instruct (temperature 0) using three prompts (appendix, prompts P1–P3): a zero-shot prompt and two prompts designed to differ in the number of worked examples, which, as retained, both include all five (appendix, supplementary methods). Combinations returned by at least two of the three were kept (self-consistency ^23^). The code of the full run, including this vote, was not retained; its output is available to editors and reviewers on request. The model returned output for 2,279 descriptions (the reason the other 217 produced none was not recorded), of which 2,261 had structured content.

#### Consensus network

Each model’s output was parsed into a drug-to-target graph, and drug–target pairs reported by at least two of the six models formed the consensus network (appendix, algorithm, steps 1–2). Network-level statistics compared names after lowercasing and collapsing whitespace; all downstream analyses used names mapped to curated references (see Entity validation), with the six graphs merged again so that different spellings of one drug or gene pooled their model support.

#### Prompt design

The extraction and validation prompts (appendix, P1–P6) give the model a role, define its entities (P1–P4 with explicit inclusion and exclusion rules), and require JSON output; only the literature and validation prompts (P4, P6) forbid inference or guessing, and the retrieval prompt (P5) is a short instruction without entity rules.

#### Entity validation

During validation of the extracted entities, we found drug classes, name variants, and partial gene-name matches; we therefore mapped drug and target names to curated references where possible and report the corrected analysis. Drug names were mapped to ChEBI (release 257) ^24^; names matching an entry with a chemical structure, or a structure-less entry without subclasses such as a therapeutic antibody, were kept as drugs, and salt and stereoisomeric forms of one compound were merged. Drug classes, labels local to one paper, chemical elements, non-drug terms, and names not found in ChEBI were set aside. Target names were mapped to HGNC gene symbols ^25^ by exact symbol or name; gene families were not mapped. Trial and RAG names were mapped with the same rules, and each drug was flagged as FDA-approved or not using Drugs@FDA (descriptive only). To measure mapping accuracy, random samples of 200 drug and 200 target names that had been mapped through a changed spelling rather than by exact match were judged by three LLMs not used for extraction (gpt-oss-120b, GLM-5.3, and Kimi-K2.7; National Research Platform; temperature 0), following our published protocol ^26^; unanimous votes were final, and for votes that were not unanimous the final judgment was made by Claude (Anthropic; claude-opus-5-5). Rules, prompt, and results are in the appendix (supplementary methods; prompt P6; Supplementary Table S4).

### Estrogen signaling pathway overlay

#### Pathway and branches

We parsed the KEGG estrogen signaling pathway (hsa04915) ^27^ into a directed graph of 63 gene nodes and 68 interactions. A breadth-first search from every gene node, up to a depth of 25 edges, produced 383 downstream signaling branches, each assigned by its genes to one or more of five signaling arms (Genomic/Nuclear, EGFR/MAPK, PI3K/AKT, PKA/cAMP, and PLC*β*/PKC) or to a residual Other category (Supplementary Table S2).

#### Drug–target–branch mapping

Each consensus drug–target pair was matched to branch genes through the target’s HGNC symbol; only exact matches were used. Each match produced a drug–target–branch triple recording the gene’s position, its arm, and the model count (appendix, algorithm, steps 3–4).

### Drug combination discovery

#### Complementary pair enumeration

To favor pairs whose drugs act on different parts of the pathway, we excluded pairs in which one drug’s branches are fully contained in the other’s: with ℬ the set of pathway branches reached by a drug’s targets, drugs A and B formed a candidate pair (compare the complementary exposure pattern ^10^) when

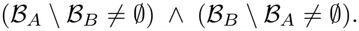

The search was restricted to the 500 drugs with the broadest branch coverage (appendix, algorithm, step 5).

#### Clinical trial verification

A candidate pair had trial precedent when both of its drugs, as mapped, co-appeared in at least one of the 2,261 trial records with structured output, excluding relations that denote comparison or sequence rather than co-administration (appendix, supplementary methods; algorithm, step 6).

### Multi-evidence ranking

#### Trial-guided RAG target identification

Using retrieval-augmented generation (RAG) ^28^, for the drugs in each trial combination five open-weight LLMs (Phi4, Aya-Expanse-32b, Gemma3-27b, Mistral-7b, and Qwen2.5-32b) extracted drug–protein pairs from the three most similar abstracts (BioBERT embeddings ^29^) in a separate corpus of 10,000 ER^+^ breast cancer abstracts (appendix, prompt P5). Pairs reported by at least two models formed the RAG consensus, which was compared with the literature extraction against the full consensus network (layer 1) and against the drug–target–branch mappings (layer 2), using Jaccard similarity and a one-sided Fisher’s exact test.

#### Ranking and evaluation

To order candidate pairs for expert review, ComboRank combined four evidence dimensions by robust rank aggregation (RRA) ^30^: pathway coverage (PC; size of the branch union), LLM consensus (LC; sum of per-drug model counts), RAG support (RS; sum of per-drug RAG model counts), and cross-check (CC; number of the pair’s drugs confirmed by both extraction methods, 0–2). Trial records were not a direct ranking input, because trial precedent is the benchmark. With fractional ranks *ρ* = rank*/N* in each dimension, a pair’s score *S*_*i*_ is the smallest beta-distribution probability across its *K* = 4 sorted fractional ranks:

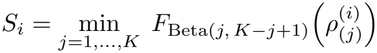

We measured how well the ordering recovered pairs with trial precedent by precision, recall, and F1 among the top *k* pairs, compared it with Borda and Dowdall aggregation, and, because RS and CC derive from RAG searches seeded with trial drugs, repeated the evaluation with PC and LC alone (appendix, supplementary methods).

### Statistical analysis

Proportions are reported with 95% CIs calculated by the Wilson score method. Agreement between the retrieval-augmented and bulk extractions was tested with a one-sided Fisher’s exact test, reported with the odds ratio, and summarized by Jaccard similarity. Pairs with and without trial precedent were compared with two-sided Mann– Whitney tests (exploratory). Ranking performance was measured by precision, recall, and F1 among the top *k* pairs; enrichment over chance is precision divided by the proportion of all pairs with trial precedent. Analyses used Python 3.13.5 with SciPy 1.18.1, pandas 3.1.1, NetworkX 3.7, and RDKit 2026.09.2.

## Results

Of 2,496 trial descriptions retrieved, Phi-3.5-mini-instruct returned output for 2,279 (the reason the other 217 produced none was not recorded), of which 2,261 were structured; 1,652 named at least one drug and 1,559 described at least one drug combination, giving 2,609 pairwise combination relationships (excluding 12 with an empty drug name) across 1,938 unique drug names (Table 1). In the literature, 587,256 of the 595,122 records had an abstract and were read by each of the six LLMs; an additional model, Llama-3.1-70B, was run but excluded because its output file was corrupt and truncated. Output quality varied across models: noisy responses ranged from 94 for Qwen-2.5-7B to 301,536 (51.3%) for Gemma-2B (Table 1), and per-model networks ranged from 438,481 edges (Gemma-2B) to 1,024,220 edges (Llama-3.1-8B). Requiring agreement between at least two models produced a consensus network of 197,336 drug names and 176,767 target names (before mapping), and 760,743 drug–target edges, 25.7% fewer edges than the largest single-model network (Table 1; Supplementary Table S1; Figure 2). Agreement was graded: of the 760,743 consensus edges, 393,344 (51.7%) were reported by two models, 181,053 by three, 101,132 by four, 58,219 by five, and 26,995 (3.5%) by all six.

**Table 1:**
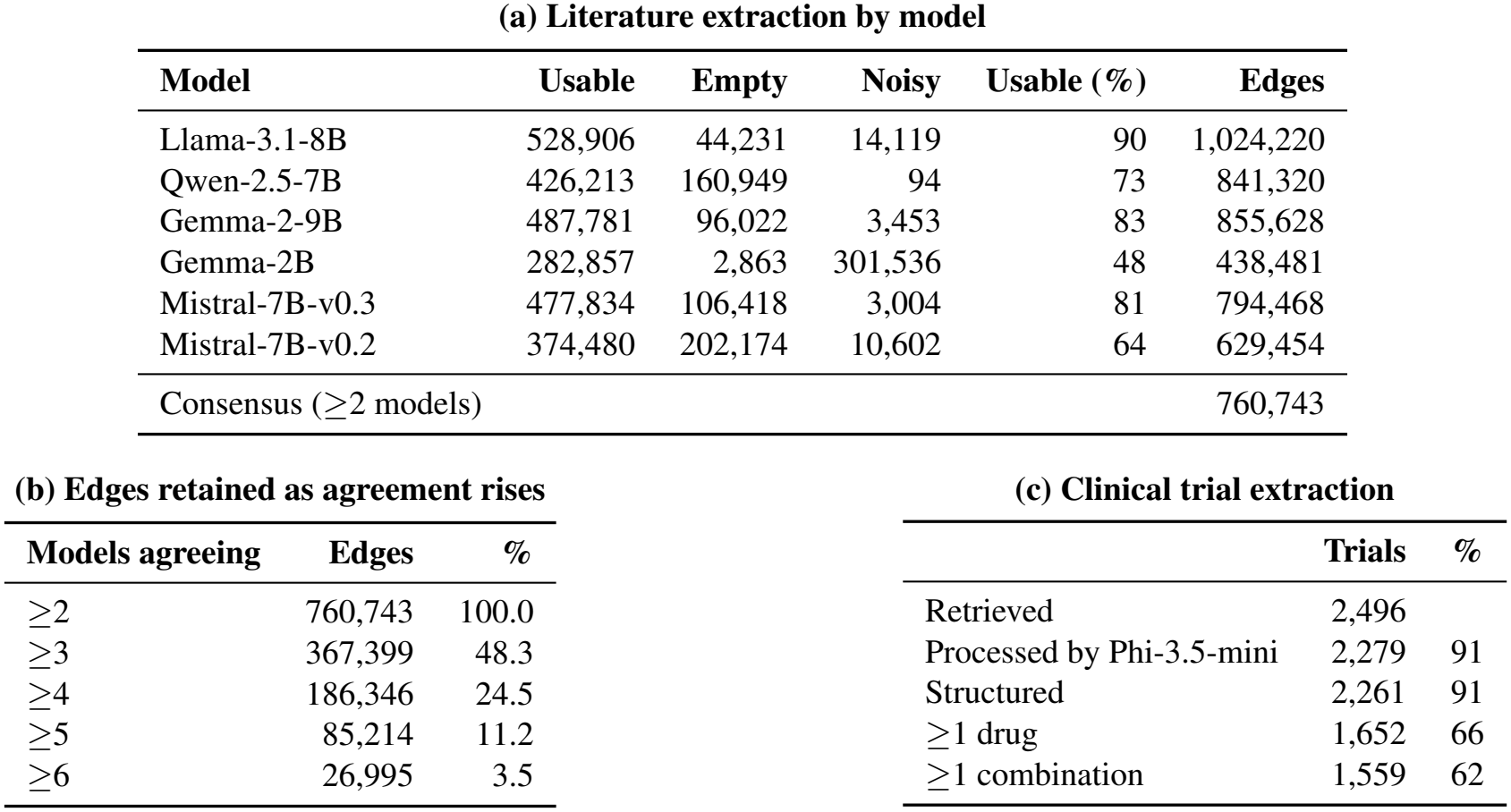
LLM-driven evidence extraction (the values shown in Figure 2). (a) Literature extraction by each of the six LLMs over 587,256 abstracts: usable, empty, and noisy responses, the usable share, and the drug–target edges in each model’s network. The consensus network keeps edges reported by at least two models. (b) Consensus edges retained as the number of agreeing models rises; percentages are of all 760,743 consensus edges. (c) Clinical trial extraction with Phi-3.5-mini-instruct (majority of three prompts); percentages are of descriptions retrieved.

| (a) Literature extraction by model |  |  |  |  |  |
| Model | Usable | Empty | Noisy | Usable (%) | Edges |
|---|---|---|---|---|---|
| Llama-3.1-8B | 528,906 | 44,231 | 14,119 | 90 | 1,024,220 |
| Qwen-2.5-7B | 426,213 | 160,949 | 94 | 73 | 841,320 |
| Gemma-2-9B | 487,781 | 96,022 | 3,453 | 83 | 855,628 |
| Gemma-2B | 282,857 | 2,863 | 301,536 | 48 | 438,481 |
| Mistral-7B-v0.3 | 477,834 | 106,418 | 3,004 | 81 | 794,468 |
| Mistral-7B-v0.2 | 374,480 | 202,174 | 10,602 | 64 | 629,454 |
| Consensus (≥2 models) |  |  |  |  | 760,743 |
| **(b) Edges retained as agreement rises** |  |  |  |  |  |
| **Models agreeing** | **Edges** | **%** |  |  |  |
| ≥2 | 760,743 | 100.0 |  |  |  |
| ≥3 | 367,399 | 48.3 |  |  |  |
| ≥4 | 186,346 | 24.5 |  |  |  |
| ≥5 | 85,214 | 11.2 |  |  |  |
| ≥6 | 26,995 | 3.5 |  |  |  |
| **(c) Clinical trial extraction** |  |  |  |  |  |
|  | **Trials** | **%** |  |  |  |
| Retrieved | 2,496 |  |  |  |  |
| Processed by Phi-3.5-mini | 2,279 | 91 |  |  |  |
| Structured | 2,261 | 91 |  |  |  |
| ≥1 drug | 1,652 | 66 |  |  |  |
| ≥1 combination | 1,559 | 62 |  |  |  |

**Figure 2:**
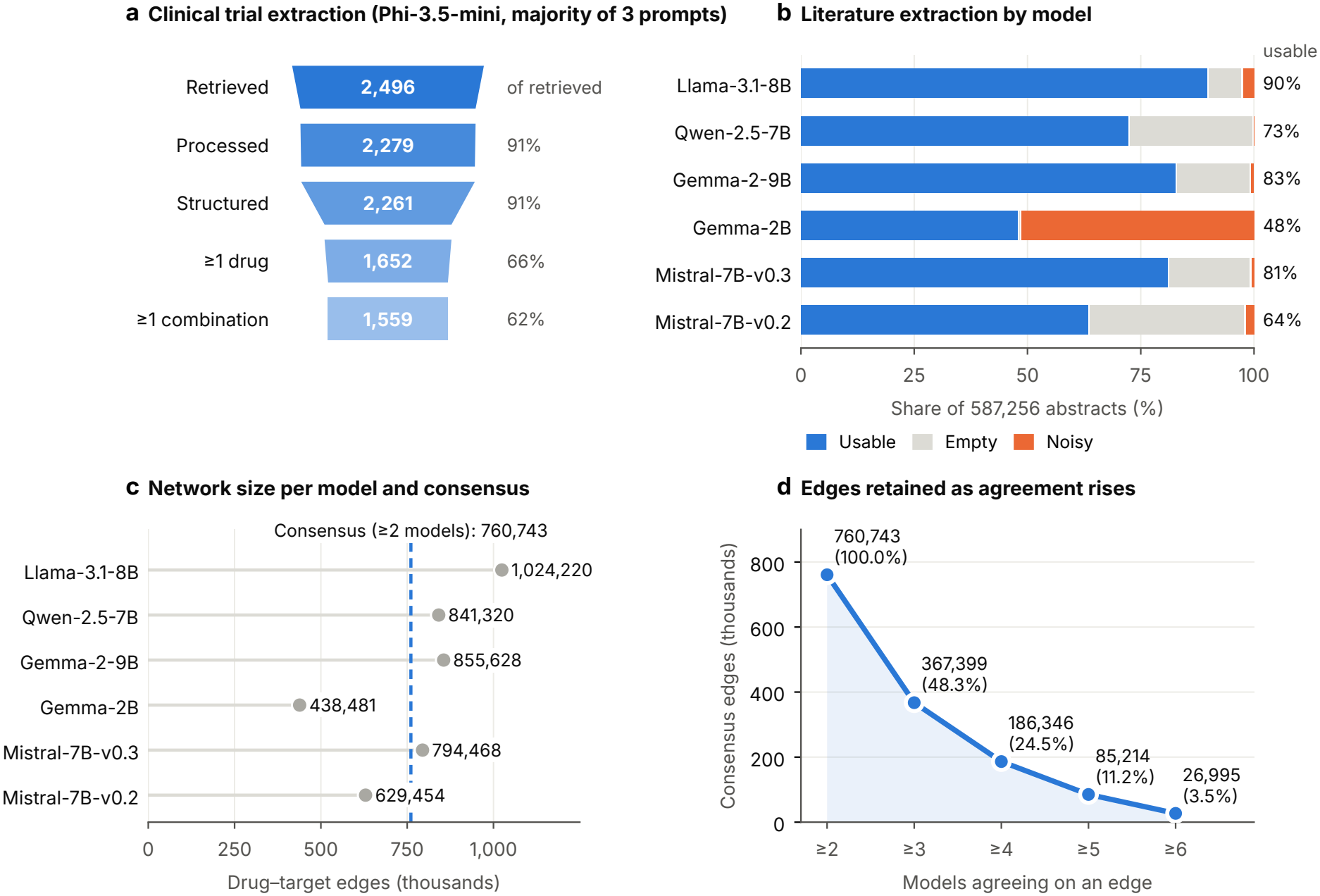
LLM-driven evidence extraction. (a) Clinical trial extraction with Phi-3.5-mini-instruct (majority of three prompts): trial descriptions retrieved, processed by Phi-3.5-mini-instruct, with structured content, naming at least one drug, and describing at least one combination; percentages are of descriptions retrieved. (b) Outcome of each of the six LLMs on the 587,256 abstracts: usable, empty, or noisy responses; percentages give the usable share. (c) Drug–target edges in each model’s network; the dashed line marks the consensus network (agreement of at least two models). (d) Consensus edges retained as the number of agreeing models rises from at least two to all six.

Of 782,470 distinct drug names in the six models’ output, 112,925 were mapped to ChEBI, corresponding to 18,065 distinct drugs; most of the rest were drug classes (270,284) or names not found in ChEBI (389,203). Of 1,002,209 target names, 100,562 were mapped to HGNC genes. In the first validation sample, 89.5% of drug and 88.0% of target mappings made through a changed spelling were judged correct; after the mapping rules were corrected for the error types found (trademark symbols, combinations, gene families, and short legacy symbols), a fresh sample gave 91.0% (95% CI 86.2–94.2) and 94.0% (89.8–96.5). Remaining errors involved lists and combination products, stereoisomers merged by structure, and contradictory source text.

Parsing the estrogen signaling pathway yielded 383 downstream branches, starting at 29 genes: 324 Genomic/Nuclear, 33 EGFR/MAPK, 16 PKA/cAMP, four PI3K/AKT, three PLC*β*/PKC, and three Other (Table 2; Supplementary Table S2). Mapping the consensus network onto these branches produced 401,435 drug–target–branch triples involving 2,187 drugs and 4,155 drug–target pairs; 35 (55.6%) of the 63 pathway genes had at least one drug, and all 383 branches were reached. The broadest coverage belonged to estradiol (365 branches), followed by dasatinib, ethanol, the phorbol ester PMA, and retinoic acid (364 each); tamoxifen reached 361 (Table 2; Figure 3). Several of the broadest compounds are endogenous hormones or laboratory reagents rather than repurposing candidates; they were retained, and each drug is flagged as FDA-approved or not.

**Table 2:**
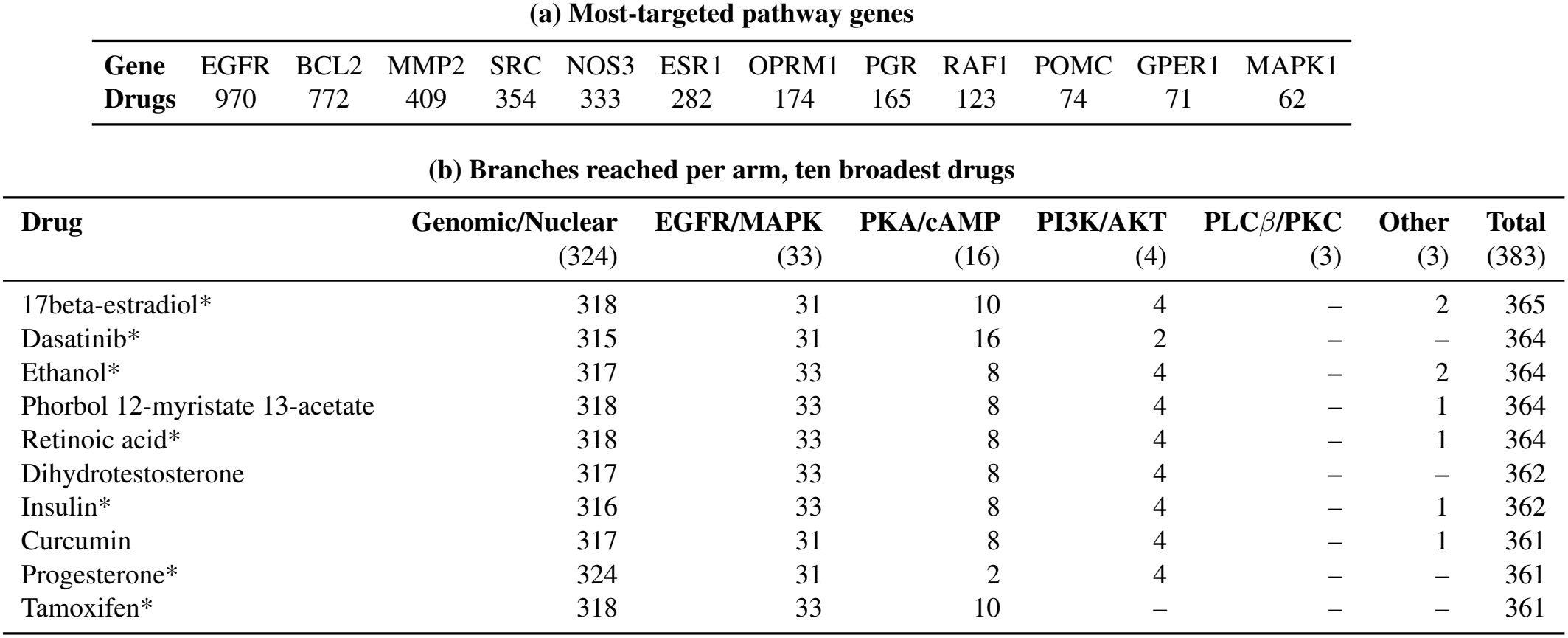
Estrogen signaling pathway overlay (the values shown in Figure 3). (a) The twelve pathway genes targeted by the most drugs (Figure 3c). (b) Branches reached in each signaling arm by the ten drugs with the broadest coverage (Figure 3b); numbers in parentheses give the branches in each arm, and drugs tied on total are listed alphabetically; * FDA-approved (Drugs@FDA).

| **(a) Most-targeted pathway genes** |  |  |  |  |  |  |  |  |  |  |  |  |
| **Gene** | EGFR | BCL2 | MMP2 | SRC | NOS3 | ESR1 | OPRM1 | PGR | RAF1 | POMC | GPER1 | MAPK1 |
|---|---|---|---|---|---|---|---|---|---|---|---|---|
| **Drugs** | 970 | 772 | 409 | 354 | 333 | 282 | 174 | 165 | 123 | 74 | 71 | 62 |
| **(b) Branches reached per arm, ten broadest drugs** |  |  |  |  |  |  |  |  |  |  |  |  |
| **Drug** | **Genomic/Nuclear**<br>(324) | **EGFR/MAPK**<br>(33) | **PKA/cAMP**<br>(16) | **PI3K/AKT**<br>(4) | **PLCβ/PKC**<br>(3) | **Other**<br>(3) | **Total**<br>(383) |  |  |  |  |  |
| 17beta-estradiol* | 318 | 31 | 10 | 4 | – | 2 | 365 |  |  |  |  |  |
| Dasatinib* | 315 | 31 | 16 | 2 | – | – | 364 |  |  |  |  |  |
| Ethanol* | 317 | 33 | 8 | 4 | – | 2 | 364 |  |  |  |  |  |
| Phorbol 12-myristate 13-acetate | 318 | 33 | 8 | 4 | – | 1 | 364 |  |  |  |  |  |
| Retinoic acid* | 318 | 33 | 8 | 4 | – | 1 | 364 |  |  |  |  |  |
| Dihydrotestosterone | 317 | 33 | 8 | 4 | – | – | 362 |  |  |  |  |  |
| Insulin* | 316 | 33 | 8 | 4 | – | 1 | 362 |  |  |  |  |  |
| Curcumin | 317 | 31 | 8 | 4 | – | 1 | 361 |  |  |  |  |  |
| Progesterone* | 324 | 31 | 2 | 4 | – | – | 361 |  |  |  |  |  |
| Tamoxifen* | 318 | 33 | 10 | – | – | – | 361 |  |  |  |  |  |

**Figure 3:**
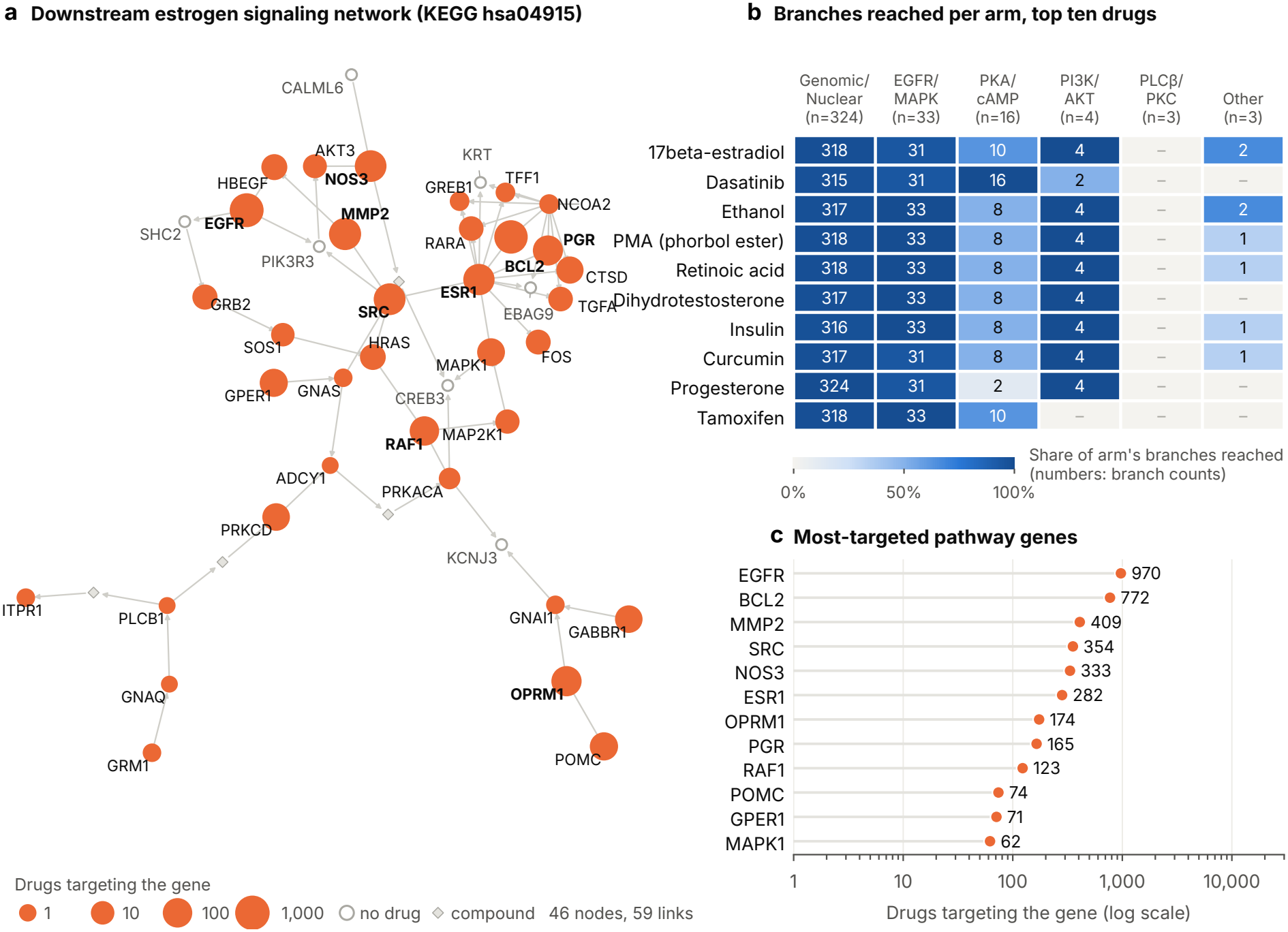
Estrogen signaling pathway overlay. (a) The downstream estrogen signaling network (KEGG hsa04915): 42 genes and four compounds on the 383 enumerated branches, with 59 links; node size shows the number of drugs targeting each gene, and hollow nodes have none. (b) Branches reached in each signaling arm by the ten drugs with the broadest coverage; shading shows the share of each arm’s branches reached, and cell numbers are branch counts. (c) The twelve pathway genes targeted by the most drugs (log scale).

Restricting the search to the 500 drugs with the broadest coverage gave 124,750 candidate pairs, of which 81,207 (65.1%) were complementary (Figure 4); 94 (0.12%) co-occurred in at least one clinical-trial record. Most pairs joined the genomic arm with the EGFR/MAPK arm (92%), and 40% reached the PI3K/AKT arm; pairs reaching PLC*β*/PKC were rare (0.7%). EGFR was a target in 85% of pairs, and the most frequent target combination, EGFR with ESR1, occurred in 57% (Table 3; Figure 4). Compared with other pairs, pairs with trial precedent had nearly the same fraction of branches shared by the two drugs (median 0.84 vs 0.83; *p* = 0.03, reflecting a small shift in distribution) but more distinct targets (median 8 vs 6; *p <* 0.0001; two-sided Mann–Whitney tests, exploratory). After name mapping, trial records contributed 945 distinct drugs and 1,758 drug pairs.

**Table 3:**
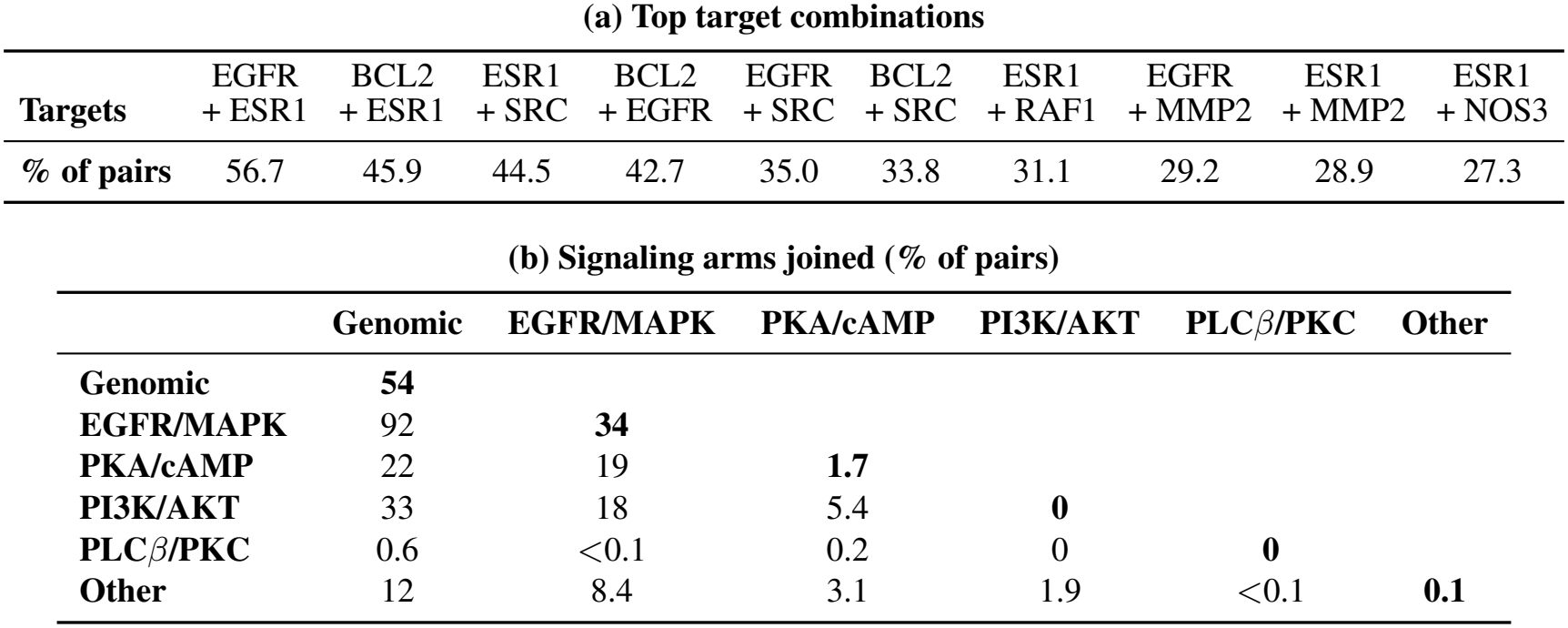
Complementary drug pairs across the estrogen signaling pathway (the values shown in Figure 4). (a) The ten target combinations seen most often across complementary pairs, one target from each drug, as a percentage of pairs; EGFR is a target in 85% of pairs. (b) Percentage of the 81,207 complementary pairs in which the branches unique to each drug fall in the given pair of signaling arms; a pair can contribute to more than one cell, and diagonal cells (bold) count pairs in which both drugs have unique branches in the same arm.

| **(a) Top target combinations** |  |  |  |  |  |  |  |  |  |  |
| **Targets** | EGFR<br>+ ESR1 | BCL2<br>+ ESR1 | ESR1<br>+ SRC | BCL2<br>+ EGFR | EGFR<br>+ SRC | BCL2<br>+ SRC | ESR1<br>+ RAF1 | EGFR<br>+ MMP2 | ESR1<br>+ MMP2 | ESR1<br>+ NOS3 |
|---|---|---|---|---|---|---|---|---|---|---|
| **% of pairs** | 56.7 | 45.9 | 44.5 | 42.7 | 35.0 | 33.8 | 31.1 | 29.2 | 28.9 | 27.3 |
| **(b) Signaling arms joined (% of pairs)** |  |  |  |  |  |  |  |  |  |  |
|  | **Genomic** | **EGFR/MAPK** | **PKA/cAMP** | **PI3K/AKT** | **PLC*β*/PKC** | **Other** |  |  |  |  |
| **Genomic** | **54** |  |  |  |  |  |  |  |  |  |
| **EGFR/MAPK** | 92 | **34** |  |  |  |  |  |  |  |  |
| **PKA/cAMP** | 22 | 19 | **1.7** |  |  |  |  |  |  |  |
| **PI3K/AKT** | 33 | 18 | 5.4 | **0** |  |  |  |  |  |  |
| **PLC*β*/PKC** | 0.6 | <0.1 | 0.2 | 0 | **0** |  |  |  |  |  |
| **Other** | 12 | 8.4 | 3.1 | 1.9 | <0.1 | **0.1** |  |  |  |  |

**Figure 4:**
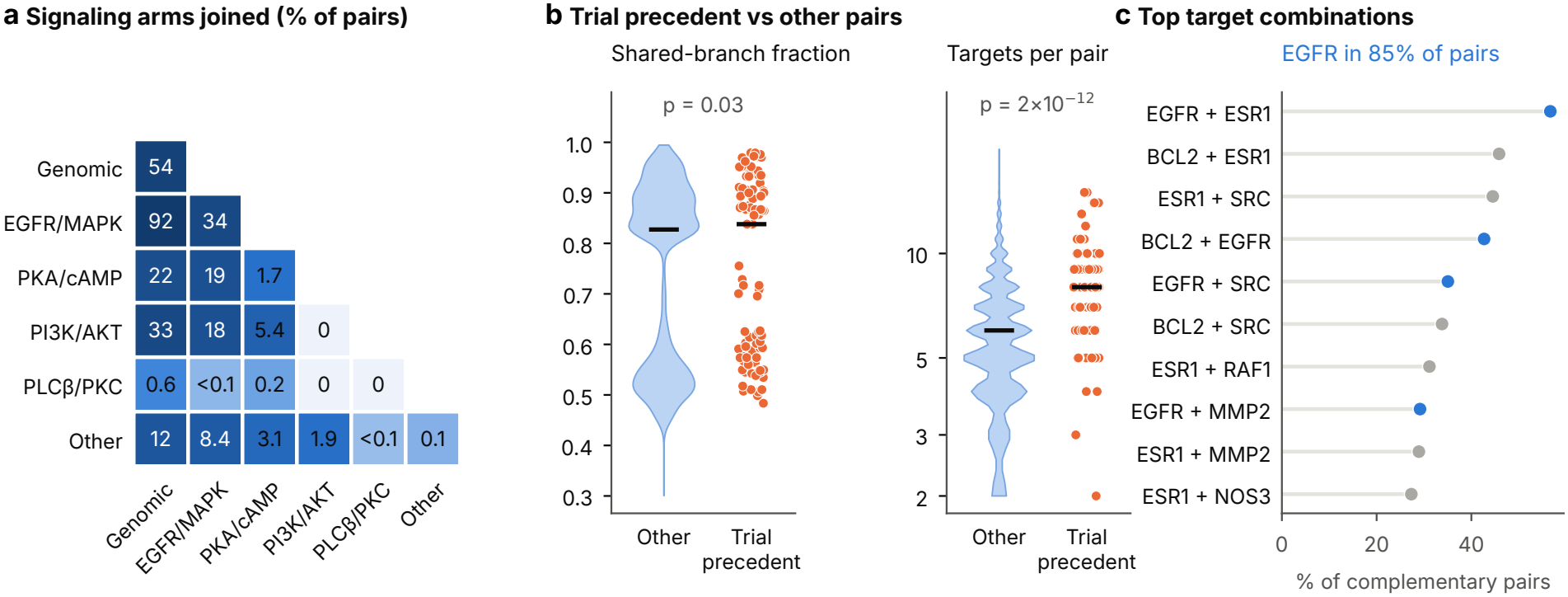
Complementary drug pairs across the estrogen signaling pathway. (a) Signaling arms joined: percentage of the 81,207 complementary pairs in which the branches unique to each drug fall in the given pair of arms; a pair can contribute to more than one cell. (b) Trial precedent vs other pairs: pairs with trial precedent (n=94, orange) compared with all other pairs (n=81,113, blue), for the fraction of branches shared by the two drugs and the number of distinct targets per pair (log scale); bars show medians; p values from two-sided Mann–Whitney tests. (c) Top target combinations: the ten target combinations seen most often across complementary pairs, one target from each drug; combinations involving EGFR are shown in blue.

Trial-guided RAG yielded 1,668 drug–protein pairs, of which 294 (217 drugs, 116 proteins) were reported by at least two models (Supplementary Table S3). Agreement with the full literature network was low (layer 1; 200 shared pairs, Jaccard 0.0005), but RAG pairs were enriched among the pathway-mapped pairs (layer 2; 32 shared pairs; one-sided Fisher’s exact test, odds ratio 10.9, *p <* 0.0001).

Without the trial-record dimension, ComboRank and Dowdall performed similarly in their top 100 pairs (precision 10.6% and 12%; ComboRank about 92 times the chance rate of 0.12%), whereas Borda gave 2% (Table 4; Figure 5). This recovery, however, came from the two RAG-based dimensions: because RAG was run on drugs from trial combinations, RAG support was present in 86% of pairs with trial precedent but in 20% of other pairs, and cross-check in 59% vs 7%. With pathway coverage and LLM consensus alone, the evidence that uses no trial information, no pair with trial precedent ranked in the top 500. The ranking is therefore a way to organize evidence for expert review, not a validated predictor of clinically tested combinations. Drug names for the top-ranked pairs are withheld pending a patent application and are available to editors and reviewers on request.

**Table 4:** Does the ranking find combinations already tested in clinical trials? (the values shown in Figure 5). All 81,207 pairs were re-ranked without the trial-record (TR) dimension, the benchmark: with pathway coverage (PC), LLM consensus (LC), RAG support (RS), and cross-check (CC); or with PC and LC only, which use no trial-derived evidence. Precision is the percentage of the top-ranked pairs with trial precedent (chance 0.12%); recall is the percentage of the 94 pairs with trial precedent found among them; best F1 is given with the cutoff at which it is reached.

|  | **Precision (%)** |  |  | **Recall (%)** |  |  |  |
|---|---|---|---|---|---|---|---|
| **Method** | Top 100 | Top 500 | Top 1,000 | Top 100 | Top 500 | Top 1,000 | **Best F1** |
| *Without TR (PC, LC, RS, CC)* |  |  |  |  |  |  |  |
| ComboRank | 10.6 | 3.3 | 2.2 | 11 | 18 | 23 | 0.12 (top 110) |
| Dowdall | 12.0 | 6.0 | 3.9 | 13 | 32 | 41 | 0.14 (top 173) |
| Borda | 2.0 | 2.5 | 1.5 | 2 | 13 | 16 | 0.05 (top 277) |
| *PC and LC only* |  |  |  |  |  |  |  |
| ComboRank | 0.0 | 0.0 | 0.2 | 0 | 0 | 2 | 0.01 (top 5,362) |
| Dowdall | 0.0 | 0.0 | 0.1 | 0 | 0 | 1 | 0.01 (top 5,362) |
| Borda | 0.0 | 0.1 | 0.2 | 0 | 0 | 3 | 0.01 (top 3,523) |

**Figure 5:**
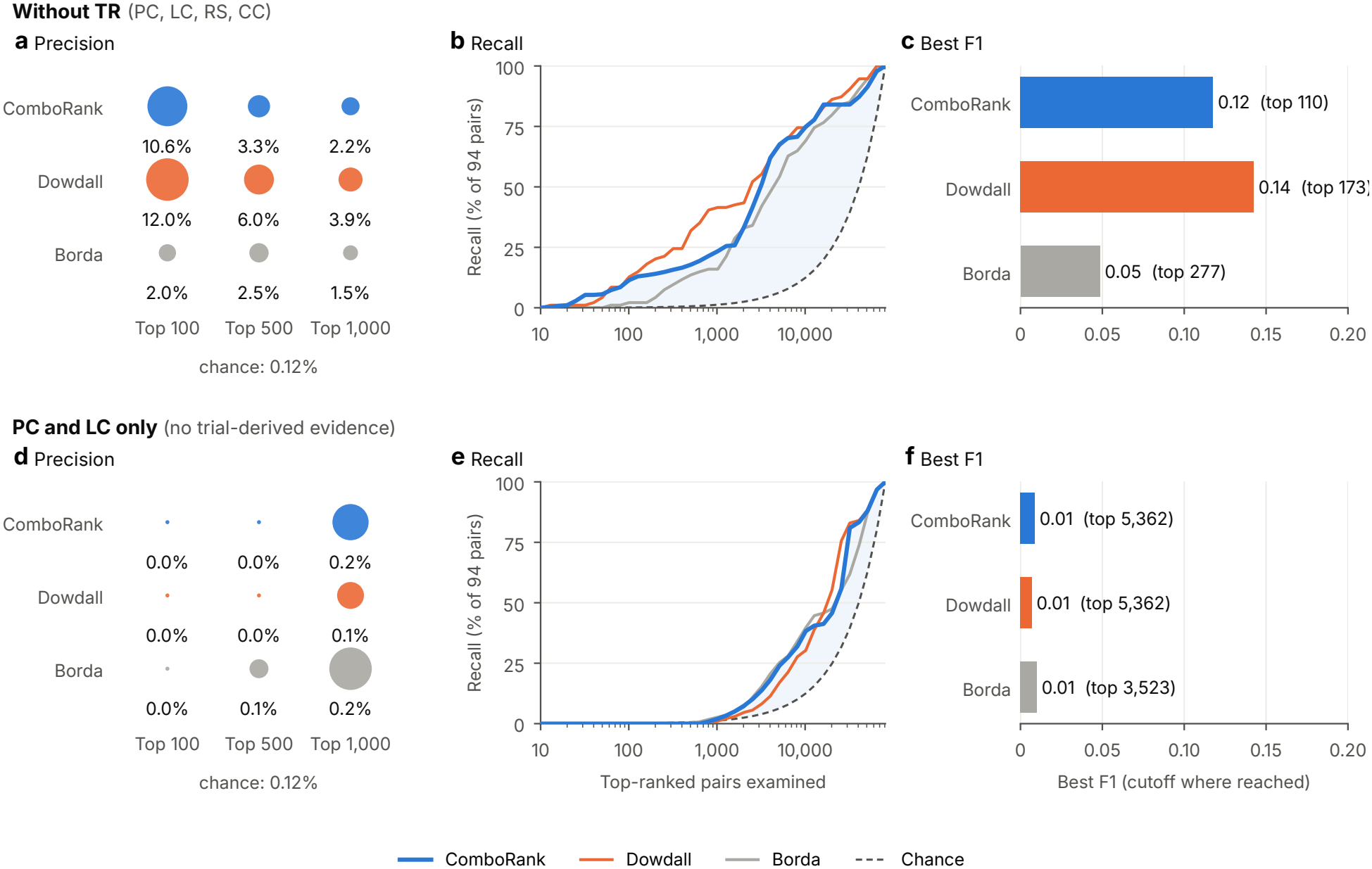
Does the ranking find combinations already tested in clinical trials? All pairs were re-ranked by ComboRank, Borda and Dowdall without the trial-record (TR) dimension, the benchmark. Top row: ranking with pathway coverage (PC), LLM consensus (LC), RAG support (RS) and cross-check (CC). Bottom row: PC and LC only, because RS and CC derive from RAG searches on trial drug names. (a, d) Precision among the top 100, 500 and 1,000 pairs; dot area is scaled within each row. (b, e) Recall: percentage of the 94 pairs with trial precedent found as the ranked list is read; dashed line, chance. (c, f) Best F1 and the cutoff at which it is reached. Chance precision is 0.12% (94 of 81,207 pairs).

## Discussion

Combinatorial drug repurposing, which pairs drugs already approved or in clinical development, offers a faster route to new treatments, but the evidence on which combinations are worth testing is scattered: publications report what each drug targets, trial records report which drugs have been given together, and curated databases organize pathway knowledge. Assembling this evidence has depended on curators reading one document at a time, a pace that cannot match tens of millions of publications and hundreds of thousands of registered trials. LLMs can read at that scale, and anchoring what they read to curated knowledge lets mechanistic evidence from the literature be weighed against clinical practice.

Speed alone, however, is not enough: LLMs can hallucinate relationships that no paper reported ^16^, and a pipeline built on such errors would only reach wrong answers faster. Our guiding principle was that no single model output should be trusted on its own. Six open-weight models from four families read every abstract independently, and a relationship was kept only when at least two reported it, so a hallucination by any one model never entered the network; trial combinations were kept only when at least two of three prompts returned them. A relationship could then shape a candidate combination only if its target was a gene of the KEGG estrogen signaling pathway. The retained relationships included well-established ones: tamoxifen and fulvestrant, approved endocrine therapies for ER^+^ breast cancer, linked to ESR1, and the EGFR inhibitor erlotinib linked to EGFR, each reported by all six models.

These safeguards reduce noise without guaranteeing truth, because several models can repeat the same misreading. Mapping names to curated references where possible, and checking samples of the mappings with a separate panel of models, exposed problems that model agreement had not: drug classes reported as drugs, one drug split into many spellings, and gene names matched by fragments (VEGFR counted as EGFR). Once the rules were corrected, more than nine in ten sampled mappings held up. For us, this is the central lesson of the study: LLMs supply the reach, and curated references and explicit validation make what they read usable.

The clearest sign that the extracted evidence is real came from two separate routes: bulk extraction by small models over every abstract, and trial-guided retrieval-augmented extraction by mostly larger models over ER^+^-focused abstracts. Before names were standardized, the two routes agreed on pathway-level drug–target pairs no more often than chance (odds ratio 1.4); afterward, the odds ratio rose to 10.9. Because only the names changed, not the models or their outputs, much of the apparent disagreement between LLM pipelines can come from how entities are named rather than from what the models read. Agreement between pipelines that differ in models, corpus, and prompts is stronger evidence than either provides alone, with the caveat that retrieval was seeded with drugs that clinicians had already chosen to test.

That caveat also limits what the ranking can claim. Its success at recovering pairs with trial precedent came only through the two retrieval-based dimensions, which carry trial information indirectly; the evidence that uses no trial information, pathway coverage and model consensus, recovered none of them near the top. Broad pathway coverage favors hub genes such as EGFR and broadly acting compounds, which is not what distinguishes combinations that reach the clinic. The ranking is best read as an ordering for expert review; a fair test of prediction would need evidence independent of trials by design, such as literature published before a set of trials was registered.

Several limitations remain. The models read abstracts only and were small to mid-sized open-weight models (2 to 32 billion parameters), chosen so that the work can be reproduced locally; larger models and full texts may extract more completely. Names not found in ChEBI were set aside, which removes some genuine drugs, and merging by structure joins stereoisomers that can differ in activity; the remaining mapping errors involve drug lists, combination products, and contradictory source text. Compounds the prompt meant to exclude, such as ethanol and the phorbol ester PMA, were retained to avoid filtering after seeing the results. Validation relied on models rather than human curators. The code of the full trial-extraction run was not retained, and two of its three prompts were nearly identical; its output is available to editors and reviewers, but the extraction itself cannot be rerun exactly. Trial records show that drugs were given together, not that the combination worked; no candidate has been tested experimentally, and the analysis covers one pathway in one disease.

The steps of this pipeline, from extraction and name mapping to validation against curated references and comparison with trials, are well suited to automation by LLM agents, the direction of our ongoing work. The broader point does not depend on the disease or the pathway: LLMs can read biomedical evidence at a scale beyond manual curation, and when each step is checked against curated knowledge, that reading becomes a resource experts can build on.

## Supporting information

Tables, Algorithms, and Prompts

## Contributors

AAH conceived the study, designed the algorithms and experiments, implemented the pipeline, carried out the analyses, and wrote the first draft of the manuscript. TEF provided pharmacological and clinical expertise and interpreted the candidate combinations. LMR provided senior scientific guidance and oversight of the research and acquired funding. All authors reviewed and edited the manuscript. AAH and TEF accessed and verified the underlying data. All authors had full access to all the data in the study and had final responsibility for the decision to submit for publication.

## Data sharing

The input data and code are publicly available at https://github.com/drahmedabdeenhamed/scaling-network-medicine-llm. The inputs are the exact PubMed query and the identifiers of the records analyzed, with a script to retrieve them (abstracts are not redistributed because some are under publisher copyright); the identifiers of the 2,496 ClinicalTrials.gov trials, with a script to retrieve their descriptions; the reference releases used for name mapping (ChEBI release 257 and HGNC, with a script to retrieve Drugs@FDA); and the prompts (appendix, P1–P6). The code covers every step of the analysis, including the reproducibility tool StatsTrace. The KEGG estrogen signaling pathway (hsa04915) is not redistributed, in line with KEGG’s terms of use; the code retrieves it. The code of the full clinical trial extraction run was not retained; its prompts and a retained prototype are provided. The set of 10,000 ER^+^ breast cancer abstracts used for retrieval-augmented generation was not retained. The analysis outputs, including the LLM extractions, the consensus network, the candidate drug pairs, and their ranking, are withheld pending a patent application; they are available only to editors and reviewers, on request to the corresponding authors, to reproduce the numbers reported.

## Acknowledgments

Ahmed Abdeen Hamed and Luis M. Rocha co-investigated the project and secured computational resource support for large-scale AI-assisted analyses. This work used resources available through the National Research Platform (NRP) at the University of California, San Diego. NRP has been developed, and is supported in part, by funding from National Science Foundation, from awards 1730158, 1540112, 1541349, 1826967, 2112167, 2100237, and 2120019, as well as additional funding from community partners.

The research reported in this publication was supported in part by the SUNY AI Platform, built on Google Cloud and funded by New York State through The State University of New York (SUNY) and The Research Foundation for SUNY (Project/Award ID: 2025-27). This work was also supported by the European Commission (Horizon Europe grant “Strategic Integration of Complex Networks and Systems for Advancing Biomedical Research”, doi:10.3030/101186558) and by Fundação para a Ciência e a Tecnologia (grant “Redundancy effects on spread and control in network dynamics: applications in computational biomedicine”, doi:10.54499/2022.09122.PTDC).

## Declaration of generative AI use

During the preparation of this work, the authors used Claude (Anthropic; claude-opus-5-5) to adapt their bioRxiv preprint (https://doi.org/10.64898/2026.09.14.728753) to this journal’s format, including drafting new sections and revising existing ones; to check the manuscript for factual, statistical, and internal consistency against the analysis outputs; to run the PubMed search reported in Research in context; to write code that standardizes names and generates the reported numbers, tables, and figures (including the reproducibility tool StatsTrace); to assemble the public code repository and the supplementary material; and to draw Figure 1. The authors also used LeapSpace (Elsevier; accessed May 25, 2026) for a literature search. The authors directed this work, reviewed and edited all output, and take full responsibility for the content of the publication. Use of LLMs in the research is described in the Methods.

## Supplementary material

The appendix contains supplementary methods, Supplementary Tables S1–S4, the workflow algorithm, and the prompt templates P1–P6. Supplementary data, released with the code, contain (a) the name-validation samples, model votes, and adjudications for both validation rounds; (b) a comparison of the analysis before and after names were standardized; (c) the records of the prior-work searches; and (d) scripts for independent checks of reported statistics. In these files, entries that name a ranked drug pair are withheld pending a patent application; the number withheld is listed in each file, and the entries are available to editors and reviewers on request. StatsTrace recomputes every number and table in the paper from the analysis outputs, which are available to editors and reviewers, and the figure scripts regenerate every figure; the outputs can be regenerated from the shared inputs and code, although the LLM extraction steps need substantial computing and may not reproduce exactly (see Data sharing).

## References

1 Cavalcante BRR, Freitas RD, Siquara da Rocha LO, Rocha GV, Gurgel Rocha CA. In silico approaches for drug repurposing in oncology: a scoping review. Frontiers in Pharmacology. 2024;15:1400029.

2 Finn RS, Martin M, Rugo HS, Jones S, Im SA, Gelmon K, et al. Palbociclib and Letrozole in Advanced Breast Cancer. New England Journal of Medicine. 2016;375(20):1925–36.

3 Burstein HJ. Systemic Therapy for Estrogen Receptor–Positive, HER2-Negative Breast Cancer. New England Journal of Medicine. 2020;383(26):2557–70.

4 Dutta B, Pusztai L, Qi Y, André F, Lazar V, Bianchini G, et al. A network-based, integrative study to identify core biological pathways that drive breast cancer clinical subtypes. British Journal of Cancer. 2012;106(6):1107–16.

5 Mejía-Pedroza RA, Espinal-Enríquez J, Hernández-Lemus E. Pathway-Based Drug Repositioning for Breast Cancer Molecular Subtypes. Frontiers in Pharmacology. 2018;9:905.

6 Goh Ki, Cusick ME, Valle DL, Vidal M, Barabási AL. The human disease network. Proceedings of the National Academy of Sciences. 2007;104(21):8685–90.

7 Sonawane AR, Weiss ST, Glass KR, Sharma AK. Network medicine in the age of biomedical big data. Frontiers in Genetics. 2019;10:294.

8 Guney E, Menche J, Vidal M, Barabási AL. Network-based in silico drug efficacy screening. Nature Communications. 2016;7:10331.

9 Rintala TJ, Ghosh A, Fortino V. Network approaches for modeling the effect of drugs and diseases. Briefings in Bioinformatics. 2022;23(4):bbac229.

10 Cheng F, Kovács IA, Barabási AL. Network-based prediction of drug combinations. Nature Communications. 2019;10:1197.

11 Conte F, Sibilio P, Fiscon G, Paci P. A Transcriptome- and Interactome-Based Analysis Identifies Repurposable Drugs for Human Breast Cancer Subtypes. Symmetry. 2022;14(11):2230.

12 Cui C, Ding X, Wang D, Chen L, Xiao F, Xu T, et al. Drug repurposing against breast cancer by integrating drug-exposure expression profiles and drug-drug links based on graph neural network. Bioinformatics. 2021;37(18):2930–7.

13 Odongo R, Demiroglu-Zergeroglu A, Ç akir T. A network-based drug prioritization and combination analysis for the MEK5/ERK5 pathway in breast cancer. BioData Mining. 2024;17(1):5.

14 Gates LE, Hamed AA. The anatomy of the SARS-CoV-2 biomedical literature: Introducing the CovidX network algorithm for drug repurposing recommendation. Journal of medical Internet research. 2020;22(8):e21169. Available from: https://www.jmir.org/2020/8/e21169.

15 Hamed AA, Fandy TE, Tkaczuk KL, Verspoor K, Lee BS. COVID-19 drug repurposing: A network-based framework for exploring biomedical literature and clinical trials for possible treatments. Pharmaceutics. 2022;14(3):567. Available from: https://www.mdpi.com/1999-4923/14/3/567.

16 Ji Z, Lee N, Frieske R, Yu T, Su D, Xu Y, et al. Survey of Hallucination in Natural Language Generation. ACM Computing Surveys. 2023;55(12):1–38.

17 Chen Q, Hu Y, Peng X, et al. Benchmarking large language models for biomedical natural language processing applications and recommendations. Nature Communications. 2025;16(3280). Available from: 10.1038/s41467-025-56989-2.

18 Babaiha NS, Rao SG, Klein J, Jacobs M, Hofmann-Apitius M. Rationalism in the face of GPT hypes: Benchmarking the output of large language models against human expert-curated biomedical knowledge graphs. Artificial Intelligence in the Life Sciences. 2024;5:100095.

19 Murali L, Gopakumar GP, Viswanathan DM, Raman R, Nedungadi PP. Integrating LLMs and Knowledge Graphs for Medical AI: Advances, Challenges, and Future Directions. IEEE Journal of Biomedical and Health Informatics. 2026.

20 Soman K, Rose PW, Morris JH, Akbas RE, Smith B, Peetoom B, et al. Biomedical knowledge graph-optimized prompt generation for large language models. Bioinformatics. 2024 09;40(9):btae560. Available from: 10.1093/bioinformatics/btae560.

21 Feng Y, Wang J, He R, Zhou L, Li Y. A retrieval-augmented knowledge mining method with deep thinking LLMs for biomedical research and clinical support. GigaScience. 2025.

22 Hamed AA, Fandy TE, Wu X. Accelerating Complex Disease Treatment Through Network Medicine and GenAI: A Case Study on Drug Repurposing for Breast Cancer. In: 2024 IEEE International Conference on Medical Artificial Intelligence (MedAI). IEEE; 2024. p. 354–9.

23 Zhou Z, Yuhao T, Li Z, Yao Y, Guo LZ, Li YF, et al. A Theoretical Study on Bridging Internal Probability and Self-Consistency for LLM Reasoning. In: Advances in Neural Information Processing Systems. vol. 38; 2025. p. 97089–122.

24 Hastings J, Owen G, Dekker A, Ennis M, Kale N, Muthukrishnan V, et al. ChEBI in 2016: Improved services and an expanding collection of metabolites. Nucleic Acids Research. 2016;44(D1):D1214–9.

25 Seal RL, Braschi B, Gray K, Jones TEM, Tweedie S, Haim-Vilmovsky L, et al. Genenames.org: the HGNC resources in 2023. Nucleic Acids Research. 2023;51(D1):D1003–9.

26 Hamed AA, Rocha LM. Protocol for evaluating ChatGPT in biomedical association generation and verification using a RAG-enabled, cross-model majority voting workflow. STAR Protocols. 2026;7:104533.

27 Kanehisa M, Furumichi M, Sato Y, Matsuura Y, Ishiguro-Watanabe M. KEGG: biological systems database as a model of the real world. Nucleic Acids Research. 2025;53(D1):D672–7.

28 Lewis P, Perez E, Piktus A, Petroni F, Karpukhin V, Goyal N, et al. Retrieval-Augmented Generation for Knowledge-Intensive NLP Tasks. In: Advances in Neural Information Processing Systems. vol. 33; 2020. p. 9459–74.

29 Lee J, Yoon W, Kim S, Kim D, Kim S, So CH, et al. BioBERT: a pre-trained biomedical language representation model for biomedical text mining. Bioinformatics. 2020;36(4):1234–40.

30 Kolde R, Laur S, Adler P, Vilo J. Robust rank aggregation for gene list integration and meta-analysis. Bioinformatics. 2012 Feb;28(4):573–80. PMID: 22247279; PMCID: PMC3278763.

