## Supplementary material for "Large language models and network medicine to identify drug combinations for estrogen receptor-positive breast cancer: a computational study": Tables, Algorithms, and Prompts

#### Supplementary methods

**Drug–target extraction.** The six models read the abstracts in concurrent batches of 500 records. Because no retrieval step was used, every abstract was presented regardless of topical relevance.

**Clinical trial corpus.** Of the 2,261 trial records with structured output, 1,652 named at least one drug and 1,559 described at least one drug combination. These records supplied clinically documented combinations and were used to check candidate pairs for trial precedent. The three combination prompts were designed to show no example, one example, and all five examples of combination language; as retained, the one-shot prompt also received all five (see Prompt design).

**Consensus network.** Each model's JSON output was parsed into a directed drug-to-target graph (NetworkX), with edges carrying the number of supporting PubMed identifiers (PMIDs), the interaction types, and the PMIDs themselves. For each drug–target pair we recorded how many models reported it and its summed frequency, and kept pairs reported by at least two models, to reduce errors specific to one model.

**Prompt design.** The prompts (P1–P6) were written to make each task narrow and checkable. P1–P4 and P6 give the model a persona with a clear standard: an expert in drug names and drug combinations for the trial prompts, a strict biomedical scientist specializing in pharmaceutical data for the literature prompt (P4), and a careful pharmacologist and molecular biologist for the validation prompt (P6). P1–P4 define their entities with explicit inclusion and exclusion rules. For drug–target extraction (P4), approved drugs, clinical trial candidates, and small molecules described as having therapeutic potential were included, and laboratory reagents, assay tools, endogenous metabolites, and radiolabels were excluded; P4 did not restrict targets or exclude drug classes, which were handled after extraction (entity validation). P4 also forbids inference ("do not infer relationships that are not explicitly stated") and asks for an empty answer when nothing qualifies, and P6 instructs the model never to guess. All six prompts require JSON output, so that responses could be parsed automatically and malformed ones discarded. The retrieval-augmented prompt (P5) is shorter: it casts the model as a biomedical research assistant, gives it the three retrieved abstracts and the drug names of one trial combination, and asks for drug–protein pairs on a signaling pathway associated with $ER^{+}$ breast cancer; it sets no inclusion or exclusion rules and does not forbid inference. The combination prompts (P1–P3) defined combination language explicitly, listing trigger phrases such as "in combination with", "combined with", and "+", and were designed to differ in how many examples of that language they showed: none (zero-shot), one (one-shot), or all five (n-shot). In the retained code, however, the example-insertion step adds all five examples to both the one-shot and the n-shot prompt, which otherwise differ only in their heading; the vote therefore combines one zero-shot and two near-identical few-shot responses, and guards less against prompt-specific artifacts than intended. The code of the full run was not retained.

**Entity validation: drug names.** Drug names were matched to ChEBI names and synonyms, ignoring case, punctuation, and spacing, and allowing for abbreviations in parentheses, plurals, and trademark symbols. A name was kept as a drug when it matched a ChEBI entry with a chemical structure, or a structure-less entry without subclasses, such as a therapeutic antibody. Names of the same compound, including salt and stereoisomeric forms, were merged

by structure (largest fragment, neutralized, first block of the InChIKey), and each drug is reported under its most frequent spelling in the literature. Names that matched a ChEBI role or class (for example, "aromatase inhibitor"), names containing class terms, labels local to one paper (for example, "compound 4"), chemical elements, and, among names not resolved by ChEBI, terms on a 640-item list of non-drug terms from our earlier work[1] (for example, "radiotherapy") were removed. Names not found in ChEBI were set aside. Each drug was flagged as FDA-approved or not using Drugs@FDA (openFDA, accessed November 11, 2026), by name; the flag was descriptive and was not used for filtering.

**Entity validation: target names.** A target name was kept only when, after removal of generic words such as "gene", "protein", or "kinase", it equaled a gene's approved symbol, previous symbol (four or more characters), or approved, previous, or alias name in HGNC. Alias symbols and shorter previous symbols (for example, "AKT", "ERK", or "ESR") were not used, because they often denote a gene family or another entity, and names denoting a gene family (for example, "Src kinases" or "EGFR family") were not mapped. Lists (for example, "MMP2 and MMP9") were mapped to every gene listed, only when every part mapped.

**Entity validation: accuracy.** Random samples of 200 drug and 200 target names that entered the pathway analysis and had been mapped through a changed spelling (for example, with an abbreviation in parentheses removed), rather than by a direct match to the reference, were judged with prompt P6. In the first sample (seed 20261111), the main error types were trademark symbols read as letters, combinations mapped to one of their drugs, gene families mapped to one gene, and short legacy gene symbols; the rules were corrected for these errors, the analysis was rerun, and a fresh sample (seed 20261112) was drawn to measure the accuracy of the corrected rules (Supplementary Table S4).

**Pathway and branches.** The KEGG pathway was parsed from KEGG Markup Language (KGML), with interactions annotated by type (gene–gene, protein–protein, protein–compound) and subtype (for example, expression, activation, binding, phosphorylation). Signaling arms were defined by their genes: Genomic/Nuclear (ESR1, NCOA, SP1), EGFR/MAPK (EGFR, KRAS, RAF, MEK, MAPK), PI3K/AKT (PIK3CA, PTEN, AKT, MTOR), PKA/cAMP (ADCY, PRKACA, CREB), and PLC$\beta$/PKC (PLCB, PRKC, DAG); branches without these genes formed the Other category (for example, GABBR1, GNAI1, OPRM1).

**Drug combination discovery.** Restricting the pair search to the 500 drugs with the broadest branch coverage kept runtime manageable. Qualifying pairs were ordered by the size of the branch union, then the size of the target union, then the combined model count.

**Clinical trial verification.** Compound names in trial records (for example, "Drug A + Drug B") were split on common separators (/, +, comma, *and*, *with*, *plus*, *or*, &) and mapped to ChEBI with the same rules as literature drugs; relations that denote comparison or sequence rather than co-administration (for example, *versus*, *compared with*, *alone*, *followed by*) were excluded. Only exact matches of mapped drugs were used. The number of trial records containing a pair was its evidence depth.

**Trial-guided RAG target identification.** The drug names of each trial combination formed the query. The 10,000-abstract corpus was exported from the PubMed web interface (10,000 records is its export limit). Noise terms (for example, "none", "not found", "not specified") were removed before name mapping. The one-sided Fisher's exact tests were computed over all unique drug–protein pairs found by either method (371,248 pairs). Because every pair in this universe is found by at least one method, the layer 1 odds ratio is not informative, and layer 1 is summarized by Jaccard similarity only.

**ComboRank and evaluation.** Within each dimension, tied pairs received their average rank. Because several dimensions take few distinct values, many pairs share ranks; we therefore did not convert scores to significance thresholds. Hits among the top $k$ pairs counted the expected number of pairs with trial precedent when a tied block straddled position $k$. Chance precision is the share of all candidate pairs with trial precedent. Borda aggregation ranks pairs by the sum of their ranks across dimensions, and Dowdall aggregation by the sum of their reciprocal ranks.

#### Supplementary tables

Table S1: Per-model drug–target network topology and consensus network summary. Average drug out-degree = edges / drug nodes; average target in-degree = edges / target nodes. The consensus row reflects the network obtained after applying a minimum two-model agreement filter (MIN_MODEL_COUNT = 2).

| Model | Drug Nodes | Target Nodes | Edges | Avg Drug Deg. | Avg Target Deg. |
|---|---|---|---|---|---|
| Llama-3.1-8B | 285,078 | 279,644 | 1,024,220 | 3.59 | 3.66 |
| Qwen-2.5-7B | 230,077 | 230,987 | 841,320 | 3.66 | 3.64 |
| Gemma-2-9B | 287,943 | 214,416 | 855,628 | 2.97 | 3.99 |
| Gemma-2B | 99,911 | 146,487 | 438,481 | 4.39 | 2.99 |
| Mistral-7B-v0.3 | 252,732 | 268,461 | 794,468 | 3.14 | 2.96 |
| Mistral-7B-v0.2 | 215,747 | 285,804 | 629,454 | 2.92 | 2.20 |
| **Consensus (MC≥2)** | **197,336** | **176,767** | **760,743** | **3.86** | **4.30** |

Table S2: Structure of the downstream estrogen signaling pathway (KEGG hsa04915) by signaling arm. Branches are the 383 downstream paths enumerated from the pathway graph (63 genes, 68 relations; Methods), each assigned to one arm. Genes are those lying on the arm's branches, and starting genes those at which its branches begin; a gene can lie on branches of more than one arm, so arm counts do not add up to the total.

| Arm | Branches | Genes | Starting genes |
|---|---|---|---|
| Genomic/Nuclear | 324 | 39 | 22 |
| EGFR/MAPK | 33 | 23 | 19 |
| PKA/cAMP | 16 | 10 | 8 |
| PI3K/AKT | 4 | 5 | 4 |
| PLC$\beta$/PKC | 3 | 4 | 3 |
| Other | 3 | 4 | 3 |
| **All arms** | **383** | **42** | **29** |

Table S3: RAG target identification and ComboRank inputs. The RAG block summarizes target identification by five LLMs from BioBERT-retrieved abstracts (drug and protein names mapped to ChEBI and HGNC), the two-model consensus, and agreement with the literature extraction (Jaccard similarity; one-sided Fisher's exact test). Raw records are before name mapping; pairs, drugs, and targets are after. The ComboRank block lists the ranked evidence; its evaluation against trial precedent is in Table 4.

| **RAG target identification** | | | | |
|---|---|---|---|---|
| **RAG model** | **Raw records** | **Unique pairs** | **Unique drugs** | **Unique targets** |
| Gemma3-27b | 3,883 | 925 | 476 | 194 |
| Mistral-7b | 6,577 | 617 | 398 | 243 |
| Aya-Expanse-32b | 4,640 | 400 | 290 | 183 |
| Phi4 | 1,155 | 140 | 102 | 74 |
| Qwen2.5-32b | 374 | 50 | 42 | 30 |
| **Consensus (≥2 models)** | – | **294** | **217** | **116** |
| Agreement, layer 1 | 200 drug–target pairs shared with the full consensus network (Jaccard 0.0005). | | | |
| Agreement, layer 2 | 32 drug–target pairs shared with the pathway-mapped pairs (Jaccard 0.0072; odds ratio 10.9; $p < 0.001$). | | | |
| **ComboRank** | | | | |
| Candidate pairs | 81,207 complementary $ER^+$ pairs ordered by Robust Rank Aggregation. | | | |
| Ranked evidence | PC pathway coverage; LC LLM consensus; RS RAG support; CC cross-check. | | | |
| Reported, not ranked | TR trial records (the evaluation benchmark). | | | |

Table S4: Validation of the name mapping. Random samples of drug and target names that had been mapped through a changed spelling (for example, an abbreviation in parentheses removed) were judged by three LLMs not used for extraction (gpt-oss-120b, GLM-5.3, Kimi-K2.7; National Research Platform; temperature 0; prompt P6), following our published protocol [2]. Unanimous votes were accepted; votes that were not unanimous, including those with a missing answer, were adjudicated by Claude (Anthropic; claude-opus-5-5). The first sample (seed 20261111) used the initial mapping rules; the rules were then corrected for the error types found, and a fresh sample (seed 20261112) was drawn. Accuracy is the share judged correct, with Wilson 95% confidence intervals.

| **Sample** | **Names** | **n** | **Unanimous** | | **Split** | **Correct** | **Accuracy, % (95% CI)** |
|---|---|---|---|---|---|---|---|
| | | | correct | incorrect | (adjudicated) | | |
| First sample | Drugs | 200 | 172 | 10 | 18 | 179 | 89.5 (84.5–93.0) |
| First sample | Targets | 200 | 146 | 18 | 36 | 176 | 88.0 (82.8–91.8) |
| Fresh sample | Drugs | 200 | 172 | 7 | 21 | 182 | 91.0 (86.2–94.2) |
| Fresh sample | Targets | 200 | 154 | 5 | 41 | 188 | 94.0 (89.8–96.5) |

#### Workflow algorithm

**Algorithm 1:** Algorithmic workflow for ER$^+$ combinatorial drug repurposing across four methodological sections: LLM-Driven Evidence Network Construction, ER$^+$ Pathway Overlay, ER$^+$ Combination Discovery, and Multi-Evidence Ranking.

**Input:** JSON files $\mathcal{J} = \{J_1, \ldots, J_n\}$; KGML pathway $K_{\mathrm{ER+}}$; NCT combinations $\mathcal{C}_{\mathrm{NCT}}$; threshold $\theta$
**Output:** $G^*$ (consensus network); $\mathcal{T}$ (overlay triples); $\mathcal{R}$ (candidate pairs); $\mathcal{V}$ (verified pairs); $\mathcal{P}^*_{\mathrm{RAG}}$ (RAG consensus); $\mathcal{R}^*$ (prioritized ranking)

*Section 1: LLM-Driven Evidence Network Construction*;

**1. Per-LLM drug–target network construction.**;
**foreach** $J_i \in \mathcal{J}$ **do**
    Parse drug–target pairs; build $G_i = (V_i, E_i)$.;

**2. Consensus drug–target network construction.**;
Map drug names to ChEBI and target names to HGNC; set aside unmapped names.;
**foreach** $(d, t)$ *observed in* $\geq 1$ *graph* $G_i$ **do**
    $s(d, t) \leftarrow |\{i : (d, t) \in E_i\}|$;
    **if** $s(d, t) \geq \theta$ **then**
        Add $(d, t, s)$ to $G^*$.;

*Section 2: ER$^+$ Pathway Overlay*;

**3. ER$^+$ pathway parsing and branch enumeration.**;
Parse $K_{\mathrm{ER+}}$; extract $\mathcal{B} = \{B_1, \ldots, B_m\}$; for each $B_j$ record gene set $\Gamma(B_j)$.;
**4. Drug–target–branch triple generation.**;
**foreach** $(d, t) \in E^*$, $B_j \in \mathcal{B}$ **do**
    **if** $t \in \Gamma(B_j)$ **then**
        Add triple $(d, t, B_j)$ to $\mathcal{T}$.;

*Section 3: ER$^+$ Combination Discovery*;

**5. Complementary drug pair enumeration.**;
For each drug $d$, let $\mathcal{B}_d = \{B_j : (d, t, B_j) \in \mathcal{T}\}$; keep the 500 drugs with the largest $|\mathcal{B}_d|$.;
**foreach** *pair* $\{d_a, d_b\}$ *of kept drugs* **do**
    **if** $\mathcal{B}_{d_a} \setminus \mathcal{B}_{d_b} \neq \emptyset$ *and* $\mathcal{B}_{d_b} \setminus \mathcal{B}_{d_a} \neq \emptyset$ **then**
        Add $\{d_a, d_b\}$ to $\mathcal{R}$.;

**6. Clinical trial verification of candidate pairs.**;
**foreach** $c \in \mathcal{R}$ **do**
    Label $c$: SUPPORTED if $c \in \mathcal{C}_{\mathrm{NCT}}$, else UNVERIFIED; add to $\mathcal{V}$.;

*Section 4: Multi-Evidence Ranking*;

**7. Trial-guided RAG target identification.**;
Retrieve top-$k$ ER$^+$-relevant abstracts per query (BioBERT cosine similarity).;
**foreach** *LLM* $m$ **do**
    Extract drug–protein pairs $P_m^{\mathrm{RAG}}$.;

$\mathcal{P}^*_{\mathrm{RAG}} \leftarrow \{(d, p) \mid |\{m : (d, p) \in P_m^{\mathrm{RAG}}\}| \geq \theta\}$; cross-validate vs. $G^*$ and ER$^+$ overlay (Jaccard + Fisher).;
**8. ComboRank: multi-evidence prioritization.**;
For each pair $(d_A, d_B)$, build $\mathbf{e} = (|\mathcal{B}_A \cup \mathcal{B}_B|,\ s_A + s_B,\ \text{RAG count},\ \text{cross-val} \in \{0\text{–}2\})$; record the NCT count separately (not ranked).;
Compute fractional ranks $\rho_{ij} = \mathrm{rank}(e_{ij})/N$; RRA score $S_i = \min_j\ F_{\mathrm{Beta}(j, K-j+1)}\left(\rho^{(i)}_{(j)}\right)$.;
Output $\mathcal{R}^*$ sorted by $S$ (ties share their average rank).;
**return** $G^*,\ \mathcal{T},\ \mathcal{R},\ \mathcal{V},\ \mathcal{P}^*_{\mathrm{RAG}},\ \mathcal{R}^*$;

#### Prompt templates

This appendix presents the prompt templates used across the three LLM-driven extraction tasks (P1–P5) and the validation of name mapping (P6), word for word as sent to the models; only code indentation was removed. Placeholders {DOCUMENT}, {SHOTS}, {pmid}, {abstract}, {context}, {query}, {cancer_type}, {name}, and {reference} are substituted at runtime with the corresponding input values; in P6, the choices in braces depend on whether a drug or a target is judged.

##### P1: Zero-Shot Drug Combination Extraction

```
# LLM Self-Consistency Exercise: Extracting Drug Entities and Relationships

## Persona:
You are an expert in drug names and the identification of drug combinations.

## Definitions:
- **Entity**: A drug name (drug).
- **Relationship**: Any of the following - "in addition to", "combination", "in
    combination with", "+", "Combined with", "together".
- **Inclusion Criteria**:
  - Verb phrases, past participle of verbs: "combine", "add".
  - "+" symbol.
  - Adverbs are desirable in relationships.
- **Exclusion Criteria**:
  - Verbs and adjectives are NOT drug entities.

## Instruction:
Extract drug entities and relationships from the text according to the given
    definitions and inclusion/exclusion criteria.


### **Input Text:**
{DOCUMENT}


### ** Expected Output:**
only drug names and the combination signal phrase or sample using the suggested
    format below.
don't provide any explanation whatsoever

### **Expected Output Format (JSON):**
```json
{
    "entities": ["DrugA", "DrugB"],
    "relationships": [
        {"entity1": "DrugA", "relation": "in combination with", "entity2": "DrugB"}
    ]
}
```

**Prompt 1:** *Zero-shot prompt (Variant 0): no trigger-keyword example is provided. Used as the baseline in the three-variant majority-vote self-consistency workflow for drug combination extraction from ClinicalTrials.gov descriptions.*
```

##### P2: One-Shot Drug Combination Extraction

````
# LLM Self-Consistency Exercise: Extracting Drug Entities and Relationships

## Persona:
You are an expert in drug names and the identification of drug combinations.

## Definitions:
- **Entity**: A drug name (drug).
- **Relationship**: Any of the following - "in addition to", "combination", "in combination with", "+", "Combined with", "together".
- **Inclusion Criteria**:
  - Verb phrases, past participle of verbs: "combine", "add".
  - "+" symbol.
  - Adverbs are desirable in relationships.
- **Exclusion Criteria**:
  - Verbs and adjectives are NOT drug entities.

## Instruction:
Extract drug entities and relationships from the text according to the given definitions and inclusion/exclusion criteria.

### **1-Shot:**
{SHOTS}


### **Input Text:**
{DOCUMENT}


### ** Expected Output:**
only drug names and the combination signal phrase or sample using the suggested format below.
don't provide any explanation whatsoever

### **Expected Output Format (JSON):**
```json
{
    "entities": ["DrugA", "DrugB"],
    "relationships": [
        {"entity1": "DrugA", "relation": "in combination with", "entity2": "DrugB"}
    ]
}
````

**Prompt 2:** *One-shot prompt (Variant 1): designed to inject one trigger-keyword example ({SHOTS}) before the input document; in the retained code, all five examples are injected, as in Variant 2.*
````

#### P3: n-Shot Drug Combination Extraction

````
# LLM Self-Consistency Exercise: Extracting Drug Entities and Relationships

## Persona:
You are an expert in drug names and the identification of drug combinations.

## Definitions:
- **Entity**: A drug name (drug).
- **Relationship**: Any of the following - "in addition to", "combination", "in combination with", "+", "Combined with", "together".
- **Inclusion Criteria**:
  - Verb phrases, past participle of verbs: "combine", "add".
  - "+" symbol.
  - Adverbs are desirable in relationships.
- **Exclusion Criteria**:
  - Verbs and adjectives are NOT drug entities.

## Instruction:
Extract drug entities and relationships from the text according to the given definitions and inclusion/exclusion criteria.

### **n-Shot:**
{SHOTS}

### **Input Text:**
{DOCUMENT}


### ** Expected Output:**
only drug names and the combination signal phrase or sample using the suggested format below.
don’t provide any explanation whatsoever

### **Expected Output Format (JSON):**
```json
{
    "entities": ["DrugA", "DrugB"],
    "relationships": [
        {"entity1": "DrugA", "relation": "in combination with", "entity2": "DrugB"}
    ]
}
````

**Prompt 3:** *n-shot prompt (Variant 2): all five trigger-keyword examples ({SHOTS}) are injected. Combinations returned by at least two of Variants 0, 1, and 2 were kept.*
````

##### P4: Drug–Target Extraction from Biomedical Literature

```
You are a strict biomedical scientist specializing in pharmaceutical data.

Input article:
  - PMID: "{pmid}"
  - Abstract: "{abstract}"

Your Goal: Extract ONLY explicit relationships between a therapeutic drug
(or potential therapeutic candidate) and its biological target.

Rules for "Drug" Classification:
  1. INCLUDE: FDA-approved drugs, clinical trial candidates, and small
     molecules explicitly described as having therapeutic potential in the text.
  2. EXCLUDE: General chemical reagents (e.g., SDS, buffers), enzymatic tools
     (e.g., "beef liver acid DNase inhibitor" used solely for assays),
     endogenous metabolites, and radio-labels unless used therapeutically.

Strict Output Rules:
  - If the text describes a chemical interaction but the agent is a standard
    reagent or tool compound and not a drug, RETURN AN EMPTY LIST.
  - Do not infer relationships that are not explicitly stated.
  - Output must be valid JSON.

JSON Schema:
{
  "pmid": "{pmid}",
  "relationships": [
    {
      "drug": "Name of the drug",
      "target": "Name of the biological target (protein, gene, receptor, etc.)",
      "interaction_type": "Brief description of the interaction
                           (e.g., inhibitor, agonist, antagonist)"
    }
  ]
}
```

**Prompt 4:** *Drug–target extraction prompt: applied to every abstract in the curated PubMed corpus (595,122 records) by each of the six open-weight LLMs. The prompt enforces strict drug classification rules and requires JSON output with no commentary.*

#### P5: Retrieval-Augmented Drug–Target Extraction

```
You are a biomedical research assistant.

Use the following PubMed abstracts to identify drug-protein interactions relevant to
     breast cancer.

Abstracts:
{context}
Query: {query}
Task:
For each drug mentioned in the query, determine whether it interacts with a protein
    that is part of
a known signaling pathway associated with {cancer_type} breast cancer.

If a valid interaction is found, return a structured list using the exact format
    below.

Output format (strict JSON):
[
  { "drug": "drug_name", "protein": "protein_name" },
  ...
]

Do not include null results.
Do not include any explanation or commentary.
Output ONLY valid JSON.
```

**Prompt 5:** *Retrieval-augmented extraction prompt, as in the code that produced the RAG results* (`code/er+_drug-protein-interaction.ipynb`)*. For each trial combination, {query} is its drug names joined by commas, {context} is the three most similar abstracts (BioBERT embeddings) from the 10,000-abstract ER*$^+$ *corpus, and {cancer_type} is "Estrogen Receptor Positive (ER+)". Sent as a single user message to each of five open-weight LLMs (Phi4, Aya-Expanse-32b, Gemma3-27b, Mistral-7b, Qwen2.5-32b) through Ollama.*

#### P6: Validation of Name Mapping

```
System:
You are a careful pharmacologist and molecular biologist. You judge
whether two names refer to the same entity. Use only well-established
knowledge; never guess. Reply with ONLY one JSON object.

User:
Name found in a biomedical abstract: "{name}"
Reference entry it was mapped to ({ChEBI | HGNC}): {reference}
Question: do the name and the reference entry refer to the same
{drug or chemical substance | human gene or its protein product}?
Salt forms, stereoisomers and spelling variants of one drug count as the
same drug; different members of a gene family count as different genes.
Return JSON: {"same": "yes" | "no" | "unsure",
              "reason": "<one sentence>"}
```

**Prompt 6:** *Validation prompt: applied to each sampled name mapping by three LLMs not used for extraction (gpt-oss-120b, GLM-5.3, Kimi-K2.7), served on the National Research Platform at temperature 0 (Supplementary Table S4). Unanimous votes were final; for votes that were not unanimous, the final judgment was made by Claude (claude-opus-5-5).*

### References

1 Hamed AA, Fandy TE, Wu X. Accelerating Complex Disease Treatment Through Network Medicine and GenAI: A Case Study on Drug Repurposing for Breast Cancer. In: 2024 IEEE International Conference on Medical Artificial Intelligence (MedAI). IEEE; 2024. p. 354-9.

2 Hamed AA, Rocha LM. Protocol for evaluating ChatGPT in biomedical association generation and verification using a RAG-enabled, cross-model majority voting workflow. STAR Protocols. 2026;7:104533.
